# Neutrophil migration in the lung is altered by alveolar collapse and stretch

**DOI:** 10.64898/2026.05.09.723927

**Authors:** Yuqing Deng, Byungjun Kang, Linzheng Shi, Chanhong Min, Kathryn Regan, Joseph K. Hall, Abdulrahman Kobayter, Neel Sajja, Kenneth R. Lutchen, J. William Boley, Jude M. Phillip, Béla Suki, Hadi T. Nia

**Author notes:** These authors contributed equally to this work (Yuqing Deng, Byungjun Kang). **Author Contributions:** Y.D., B.S., and H.T.N. designed research; Y.D. and B.K. performed research; L.S., C.M., K.R., J.K.H., A.K., and J.M.P. contributed new reagents/analytic tools; Y.D. and N.S. analyzed data; K.R.L. contributed to data interpretation and statistical analysis; W.B. contributed new tools for the experiment design; and Y.D., B.K., L.S., C.M., K.R., J.K.H., B.S., and H.T.N. wrote the paper. **Competing Interest Statement:** The authors declare no competing interest.

## Abstract

**Rationale:** Heterogeneous alveolar collapse is prevalent in inflammatory lung conditions such as chronic obstructive pulmonary disease, acute respiratory distress syndrome, and pneumonia. Although neutrophil-released proteases contribute to the tissue remodeling that leads to alveolar collapse, how this altered mechanical environment in turn affects neutrophil migration remains largely unexplored.

**Objectives:** In this study, we investigate how alveolar collapse and stretch influence neutrophil migration and identify the mechanical and biochemical factors that govern regional migration differences.

**Methods:** We developed a novel precision-cut lung slice platform that generates collapsed vs non-collapsed regions within the same slice. Neutrophils in both regions were longitudinally imaged for up to 5 hours to quantify motility behavior. Migration mechanisms were probed using migration-related inhibitors, collagenase, and cigarette smoke extract. A crystal ribcage system, which preserves intact alveolar shape and the air-liquid interface, was also used to assess the effects of ventilation on neutrophil migration.

**Results:** Neutrophil migration was faster in the collapsed region compared to not-collapsed regions. This regional difference was eliminated by Rho-associated protein kinase (ROCK) inhibition, which selectively increased migration speed in the non-collapsed region. The regional difference persisted with the addition of collagenase and cigarette smoke extract, both of which significantly increased the migration speed in both regions. In the crystal ribcage, the preserved air-liquid interface and ventilation together enhanced neutrophil migration compared with a collapsed lung.

**Conclusions:** Alveolar collapse and stretch facilitate neutrophil migration, indicating the role of localized tissue remodeling in driving neutrophil activity and further disease progression.

## Introduction

Neutrophils are the first responders of the immune system in the lung. In pulmonary diseases or as a result of inflammatory stimulation, neutrophils are recruited to the lung where they release proteases such as collagenase and elastase that degrade the extracellular matrix (ECM) ^1^. Dysregulation of neutrophil activities are commonly observed in chronic obstructive pulmonary disease (COPD) ^2,3^, where exacerbations have a high mortality rate, and are also associated with neutrophilic inflammation ^4,5^.

Neutrophil migration is a critical indicator of neutrophil activity and function ^6^. Previous studies demonstrated that neutrophil activation and migration can be regulated by mechanical cues such as substrate stiffness and mechanical deformation ^7,8^. These findings are particularly relevant in the lung, where disease progression is characterized by altered tissue mechanics and substantial ECM remodeling. For example, in COPD, a disease predominantly caused by cigarette smoking, the presence of enlarged airspaces surrounded by collapsed alveolar regions creates a heterogeneous ECM structure with both stretched and collapsed tissue ^9^. This alveolar collapse is also observed in other pulmonary conditions such as pneumonia and acute respiratory distress syndrome ^10^. Furthermore, COPD is associated with elevated levels of proteases that actively degrade the ECM ^11^. Despite the established role of mechanical factors in regulating neutrophil behavior and the observation of dramatic mechanical heterogeneity in the emphysematous lung, how neutrophils migrate in locally stretched and collapsed regions in lung tissue, and in the presence of stimulatory compounds (e.g. cigarette smoke) and proteases remains poorly understood. Decoupling the role of mechanical cues and chemical stimuli underlying neutrophil activity in conditions such as collapsed versus stretched alveoli remains poorly understood, mainly due to the lack of appropriate model systems to mechanistically study these effects.

Precision-cut lung slices (PCLS) are a valuable tool in preclinical and translational respiratory research ^12,13^. This ex vivo model enables high-throughput studies in the native ECM with proper cellular diversity well-suited for live imaging of cellular dynamics, and hence studying disease pathology ^14^. For example, stem cell motility after lung injury was captured through longitudinal time-lapse imaging of PCLS and validated in vivo ^15^. Obernolte et al. exposed human PCLS to cigarette smoke condensate to examine inflammatory and metabolic responses ^16^. Despite these advances, however, it remains unexplored how recruited neutrophils migrate within the PCLS under conditions that mimic the mechanical and chemical (e.g., presence of cigarette smoke extract (CSE) and collagenase) microenvironment of the emphysematous lung tissue. Conventional PCLS are prepared by filling alveolar spaces with agarose, a process which holds the entire slice in a uniformly inflated state and makes it impossible to compare cell migration in collapsed and stretched regions within a single PCLS.

Here, we report a novel PCLS preparation in which thermo-reversible gelatin infiltration, combined with custom 3D-printed pin supports, generates co-existing collapsed and stretched regions within a single slice. This method allows neutrophil migration to be compared across mechanically distinct microenvironments within the same tissue preparation, eliminating between-sample variability and enabling multiple pharmacological conditions to be screened in parallel. Furthermore, we exposed PCLS to CSE and bacterial collagenase, mimicking and investigating neutrophil migration in the biochemical environment present in emphysema. We hypothesized that both the mechanical state of the ECM (stretched vs. collapsed) and exposure to emphysema-associated agents would alter neutrophil migration. We then tested whether the native alveolar shape, air-liquid interface and dynamic stretching during ventilation alter migration speed in intact lungs using the recently developed crystal ribcage system ^17–20^.

## Materials and Methods

### Mouse model

Male mT/mG mice (B6.129(Cg)-Gt(ROSA)26Sortm4(ACTB-tdTomato,-EGFP)Luo/J; JAX stock #007676, Jackson Laboratory) aged 16–20 weeks were used for all experiments. All experiments conformed to ethical principles and guidelines under protocols approved by the Boston University Institutional Animal Care and Use Committee.

To recruit neutrophils to the lung, mice received an intraperitoneal injection of lipopolysaccharide (LPS) 7 hours prior to euthanasia ^21^ and a retro-orbital injection of Ly-6G antibody 3.5 hours prior to euthanasia (Supplementary Table S1; Supplementary Fig. S2).

### PCLS preparation

After euthanasia, the chest cavity was opened to allow lung collapse. Two 24G catheters were inserted through the trachea into the left and right mainstem bronchus and ligated in place (Fig. 1a; Supplementary Fig. S2). Using syringe pumps, one lobe was inflated with 37°C agarose and the contralateral lobe with 37°C gelatin at 0.02 ml/s, with the lobe-to-gel assignment alternated across experiments. The surface of the lung was flushed with chilled Hank’s Balanced Salt Solution (HBSS) to solidify the agarose/gelatin within the alveolar spaces. One agarose-inflated lobe and one gelatin-inflated lobe were carefully dissected into chilled HBSS.

**Figure 1.**
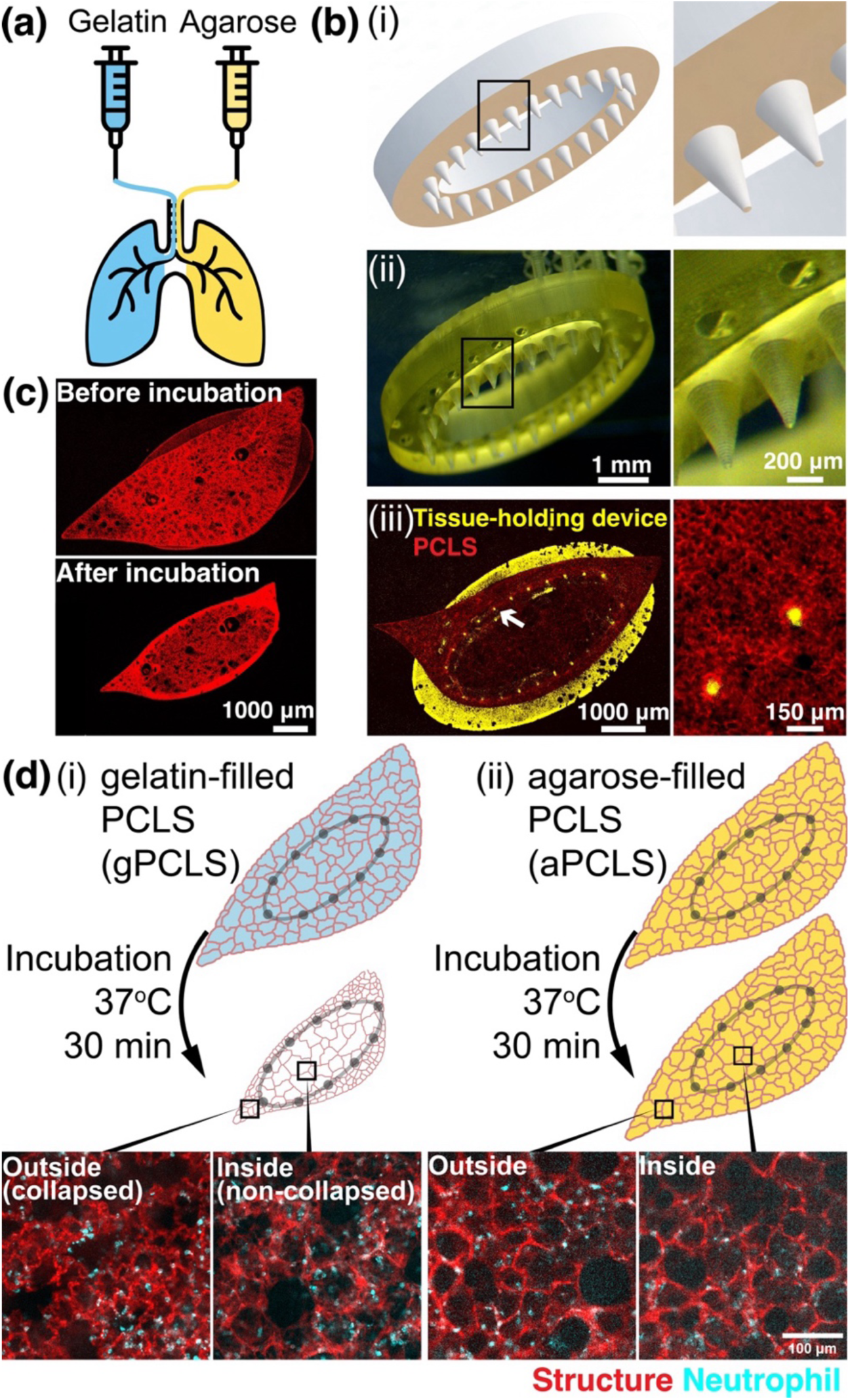
Experimental setup. (a) PCLS were prepared by simultaneous lobar filling with liquid: either 1.5% agarose or 6% gelatin at 37°C. (b) The (i) schematic and (ii) photography of a 3D-printed tissue-holding device consisting of a ring with sharp pins pointing outward with magnified insets shown at right. The zoom-in of the pins inside black box region is on the right. (iii) Pins were placed on top of the PCLS. White arrow: autofluorescence signal from the pin on top of the PCLS. (c) gPCLS collapses during incubation at 37°C as gelatin melts. (d) (i) gPCLS showed collapsed and non-collapsed region inside and outside the device. (ii) aPCLS showed same non-collapsed structure inside and outside the device.

250 μm-thick agarose-filled PCLS (aPCLS) and gelatin-filled (gPCLS) were then generated from the respective lobes using a Compresstome VF-300-0Z vibrating microtome (Precisionary Instruments). Individual PCLSs were positioned with custom 3D-printed tissue-holding device (Fig. 1b) that mechanically supported a region of the slice.

### Live imaging and quantification of neutrophil migration

PCLS were imaged on a Nikon CSU-W1 SoRa spinning disk confocal microscope in a stage-top incubation chamber (37 °C, 5% CO₂), using 640 nm excitation for Alexa Fluor® 647-labeled neutrophils (Ly-6G⁺) and 516 nm excitation for the mT/mG membrane signal (Supplementary Fig. S2). Following two-step image stabilization, neutrophils were detected and tracked in Imaris (v9.8.2, Oxford Instruments).

The mean squared displacement (MSD) was computed based on the cell tracking trajectory (Fig. 2d, i) as

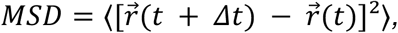

where 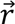 is neutrophil position at time t, and <…> brackets represent time averaging over all valid displacement pairs separated by 𝛥𝑡 across all tracked neutrophils. In anomalous diffusion, the MSD is described as

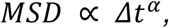

where 𝛼 is the MSD exponent calculated from the slope of a log-log linear fit over 𝛥𝑡 between 1 to 30 min (Fig. 2d, ii).

**Figure 2.**
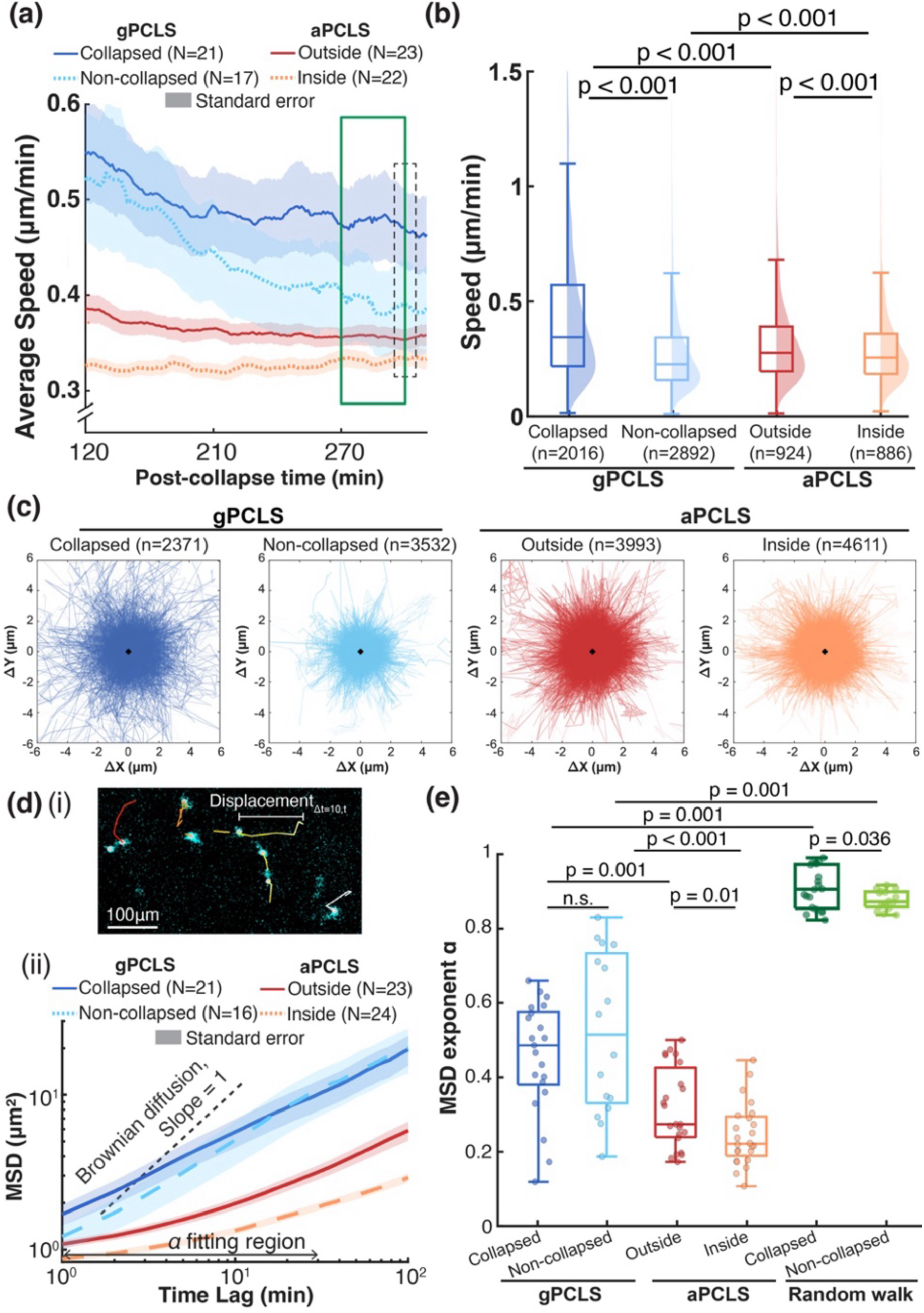
Comparison of neutrophil migration in aPCLS and gPCLS reveals higher speed, increased spreading, and reduced confinement in gPCLS. (a) Average neutrophil migration speed across different conditions between post-collapse time 150-300 min. Post-collapse time 0 was annotated in Supplementary Figure S2 as t_0_. Black dashed box (295-305 min): time window over which cell migration speeds were averaged and used for later statistical analysis of speed distribution in (b). Green box (270-300 minutes): time window used for trajectory plots in (c). (d) MSD analysis of neutrophil migration. (i) Representative image showing neutrophil (blue signal) tracks over a 10-minute-long window. Track colors are based on trace position. Displacement_Δt=10,t_ is the displacement of one neutrophil at time t with time lag Δt of 10 min. (ii) MSD as a function of time lag for neutrophils migrating in aPCLS and gPCLS. The shaded gray region indicates the fitting window (1–30 min) used to calculate the MSD exponent α. The black dashed line (slope = 1) represents the reference for Brownian diffusion. Conditions with slope below this line represent subdiffusive migration. (e) Comparison of the MSD exponent α across all regions of interest (ROI) in gPCLS, aPCLS, and random walk simulation. Each dot represents the α value from one ROI. Boxplot: center line, median; box edges, 25th and 75th percentiles; whiskers, 1.5× interquartile range; dots, outliers. N: number of ROI; n: number of cell tracking. Mice = 7 for gPCLS. Mice = 3 for aPCLS.

### Imaging of neutrophil in ex vivo lung inside crystal ribcage ^17^

After euthanasia, the lung was perfused with DMEM/F12 at 20 µL/min for at least 10 minutes, after which the lung-heart block was transferred to the crystal ribcage (Fig. 6a). Neutrophils were imaged at 37 °C under collapsed or ventilated conditions using an upright spinning-disk confocal microscope (Nikon Crest X-Light V3) at 30-s intervals. For ventilated imaging, ventilation was performed between 5-10 cmH₂O and paused for 5 s at 14 cmH₂O during each image acquisition.

### Statistics

A two-sided Wilcoxon rank sum test was applied to compare migration speed of individual cells between different condition. The p value of migration α between groups was computed using two-sample t-test.

*Additional methodologic details are provided in the Supplemental Methods*.

## Results

### A novel model generates collapsed and non-collapsed region within a single PCLS

To investigate how the alveolar stretch affects neutrophil migration, we developed a novel model that generates both collapsed and non-collapsed regions within the same precision cut lung slice (PCLS), enabling high-throughput live imaging of cell motility (Fig. 1). To compare the current method with traditional PCLS model, a conventional agarose-filled PCLS (aPCLS) were prepared simultaneously with the recently developed gelatin-filled PCLS (gPCLS) through lobar inflation with agarose and gelatin into opposite lobes of the same lung (Fig. 1a).

For each PCLS, a customized tissue-holding device (Fig. 1b, i and ii) was printed via ultra-high resolution 3D printer, and placed on top to mechanically supports a defined region via a circle of pins (Fig. 1b, iii). In gPCLS, upon incubation at 37°C, the gelatin used to inflate and prepare gPCLS melts, causing the unsupported tissue outside the device to spontaneously collapse through elastic recoil (Supplementary Video 1), while the pin-supported region maintains its inflated, stretched alveolar structure (Fig. 1c and Fig. 1d, i). This creates two spatially distinct ECM microenvironments within a single slice: a stretched region that maintains its stretched alveolar structure and a collapsed region that recapitulates structural features in locally collapsed lung, common in advanced pulmonary pathologies such as cancer, emphysema, fibrosis, and pneumonia (Fig. 1d, i).

In conventional aPCLS, solidified agarose remains within the alveolar spaces, preserving the inflated structure and precluding observation of cell behavior in mechanically collapsed environments (Fig. 1d, ii) ^12–14^. Compared to the gel-free gPCLS, the solid hydrogel itself within aPCLS may affect neutrophil migration. Using a LPS-induced inflammation model, we imaged the migration patterns of recruited and fluorescently labelled neutrophils in both gPCLS and aPCLS (Fig. 1d, blue signal; Supplementary Video 2). Since inflammation was induced systemically via intraperitoneal (IP) injection, generating aPCLS and gPCLS from the same lung ensures matched inflammation levels between the two types of PCLS.

### Alveolar collapse leads to higher neutrophil motility

Once the gelatin was removed from the gPCLS by melting at 37°C, PCLS obtained from gelatin-inflated lobes have no gelatin-filling within the alveolar spaces. At early timepoints, neutrophils in non-collapsed regions displayed the same migration speeds than in the collapsed regions (Fig. 2a, blue curves), indicating similar neutrophil recruitment and activation across regions. After 5 hours, the migration speed decreased gradually (Fig. 2a), possibly due to cellular apoptosis or NETosis-related cell death as observed in other in vitro neutrophil studies ^22^. Importantly, by 5 hours post-collapse, the 2 average speed curves diverge, resulting in faster neutrophils migrating in the collapsed regions ([median±se] gelatin: 0.34±0.01 µm/min) than non-collapsed region (0.23±0.005 µm/min) at later timepoints (Fig. 2b). This difference happens following prolonged neutrophil exposure to different stretched states of the lung ECM, possibly reflecting an adaptive switch in migration strategy ^23,24^. Trajectory plot also showed neutrophils in the collapsed regions exhibiting higher spreading of migration compared to non-collapsed region in gPCLS (Fig. 2c).

To evaluate whether these region-dependent differences could also be detected in conventional lung slices ^12,13^, we compared neutrophil migration between two regions within aPCLS, where the alveolar spaces remain both filled with a solid hydrogel. Although a difference in migration speed was also observed between regions inside and outside the tissue-holding device in aPCLS, this can be due to the initial inhomogeneous lobe inflation (Supplementary Fig. S1) or neutrophil activation from the pleural surface (typically outside the tissue-holding pins) toward the inner regions of the lobe ^25–27^, as also supported by the observed difference in speeds at early timepoints in the aPCLS (Fig. 2a). Nevertheless, the spreading of migration in aPCLS remained comparable between inside and outside regions (Fig. 2c). Besides, the neutrophil migration in aPCLS was slower compared to the migration in gPCLS (Fig. 2a), indicating that solid agarose-filling of the alveolar spaces impedes neutrophil migration.

To further characterize the nature of cell motility, we examined the mean squared displacement (MSD), which quantifies how far neutrophils spread on average from their starting point over a given time interval ^7,28^. The MSD exponent α, calculated from the MSD plot (Fig. 2d), was less than 1 in both aPCLS and gPCLS, representing a subdiffusive migration pattern (Fig. 2e), suggesting that neutrophil migration is hindered by physical and biochemical interactions between neutrophils and the surrounding ECM ^29,30^. The presence of solid agarose within the alveolar space appears to introduce additional steric and mechanical barriers to neutrophil migration, resulting in a significantly lower α compared to gPCLS (p ≤ 0.001, Fig. 2e), in which there was no solid gelatin in the alveoli. This suggests that PCLS from agarose-filled lungs have inherent limitations that prevent the widely used aPCLS from being an ideal platform to study cell migration. Overall, the migration pattern remains subdiffusive, indicating that neutrophils tend to bind to the lung ECM.

While it is experimentally difficult to isolate the contributions of tissue geometry and biological interactions to collapse-sensitive migration, we addressed this question computationally by applying a Monte Carlo random walk model that simulates neutrophil migration in PCLS under different substrate tissue structures (Supplementary Fig. S3). The random walk MSD exponent α < 1 (Fig. 2e) indicates that geometry alone determined the subdiffusive migration behavior. The simulated α for collapsed and non-collapsed regions are significantly different (p = 0.04; 0.91 ± 0.06 vs. 0.88 ± 0.03, mean ± std), suggesting that tissue geometry affects cell migration. However, the effect on collapsed geometry only showed a slightly higher α by 3%, suggesting that geometric effects, while detectable, are minor. A greater variance was observed in the experimental α (p < 0.001, F-test), likely reflecting additional biological interactions within PCLS, such as cytokine signaling, that affect α ^12,13^ and may obscure the contribution of geometry. Furthermore, experimental α values were approximately 50% lower than those from the random walk simulation (p < 0.001), indicating that binding and interactions between neutrophils and the parenchymal ECM substantially constrain migration relative to a purely geometry-driven random walk. Thus, the random walk simulations revealed that tissue geometry contributes to, but does not dominate, neutrophil migration behavior in PCLS.

### Neutrophil exhibited a distinct motility pattern in collapsed region of gPCLS

Beyond single-feature comparisons, we sought to identify higher-order motility patterns and their modulation by PCLS conditions. We quantified 93 motility features (Supplementary Table S2) capturing neutrophil movement in magnitude, spatial–temporal organization, signal characteristics, correlation structure, and entropy, and constructed a single-cell motility space to identify eight distinct motility clusters (MCs) (Fig. 3a) ^31,32^. The existence of these MCs can explain the large variance of speeds under each condition. MCs were ordered by increasing speed and further distinguished by additional features, with MC0 representing slow, tortuous movement and MC7 representing fast, directed motility (Fig. 3c). Consistent with the speed analysis in Fig. 2, a large overlap of motility space distribution was found between the two regions within aPCLS, confirming their similarity in neutrophil motility (Fig. 3b). Based on this framework, the collapsed regions in gPCLS exhibited a distinct MC distribution, such as high occurrence of MC7 (Fig. 3d), which is characterized by high average speed and progressivity (Fig. 3c). This greater proportion of neutrophils within MC7 is also consistent with the high spreading and speed identified in Fig 2. Correlation analysis further supported this pattern: the highest similarity occurred between the two aPCLS regions, whereas in gPCLS, the collapsed region showed lower correlation with all other three conditions (Fig. 3e). Taken together, the multidimensional analysis reinforces the conclusion that alveolar collapse drives a distinct, fast migratory pattern in gPCLS. And this observation is masked by the physical constraints of solid hydrogel in conventional aPCLS.

**Figure 3.**
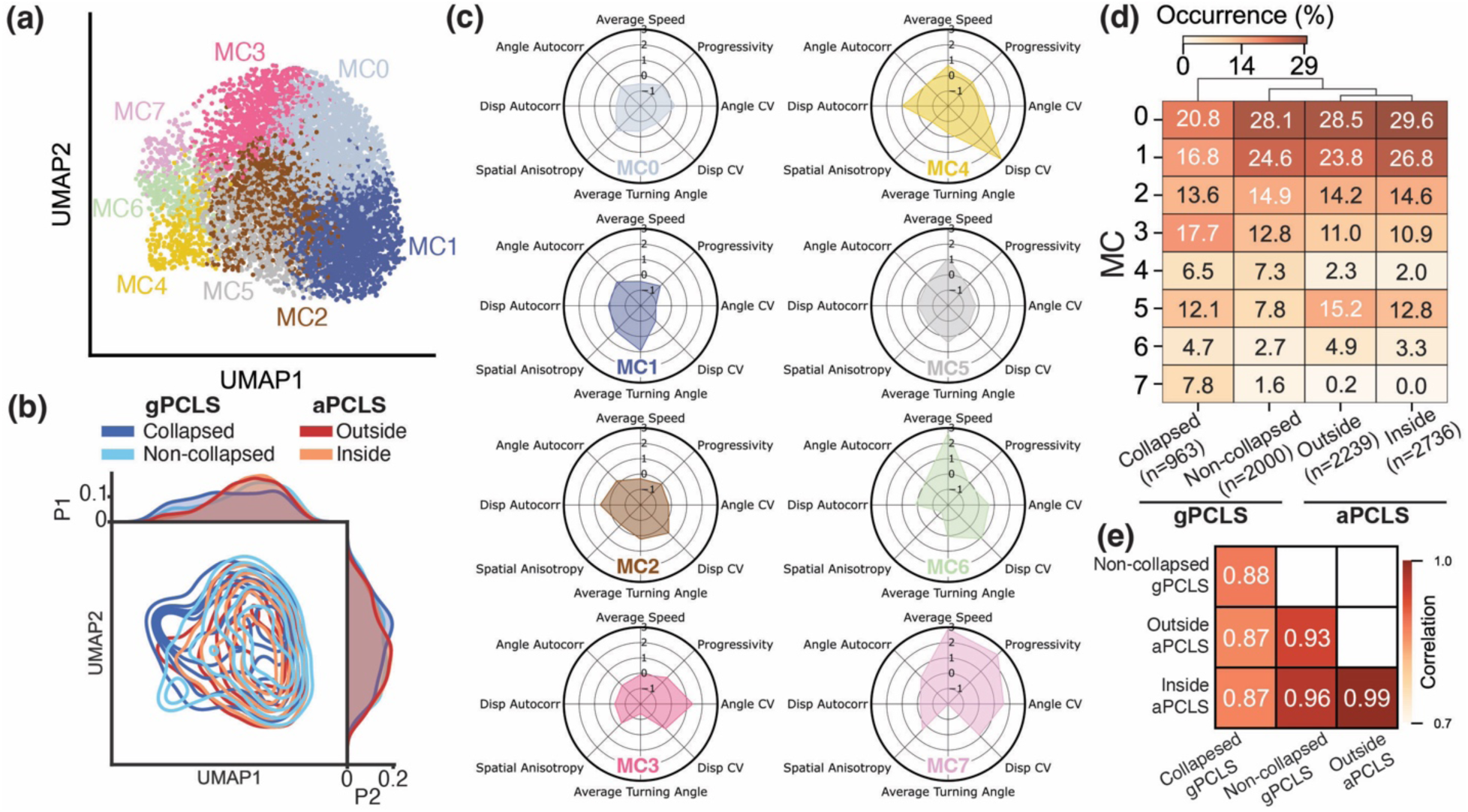
Multidimensional motility analysis of neutrophil migration in a- and gPCLS showed similar migration profile within aPCLS and differences between 2 regions in gPCLS. (a) Cluster analysis of the migration of 7938 neutrophil tracks in PCLS identified 8 distinct MCs that characterize distinct movement patterns. (b) Motility space distribution across 4 PCLS conditions. Probability distribution in UMAP1 (P1) and UMAP2 (P2) are projected to the top and right, respectively. (c) Characteristic motility features in each MC. (d) Heatmap of MC occurrence between different regions in aPCLS and gPCLS. N: number of cell tracks analysis in each PCLS conditions. (e) Pair-wise correlation map indicating similarities in the neutrophil MC distribution between PCLS conditions.

### Cell contractility mediates the alveolar collapse-dependent neutrophil migration

To further investigate the pathways regulating the collapse-sensitivity of neutrophil migration, we employed migration-related inhibitors including mechanosensitive ion channels and actomyosin contractility to probe which pathways play an important role in the speed difference between collapsed and non-collapsed regions (Fig. 4).

**Figure 4.**
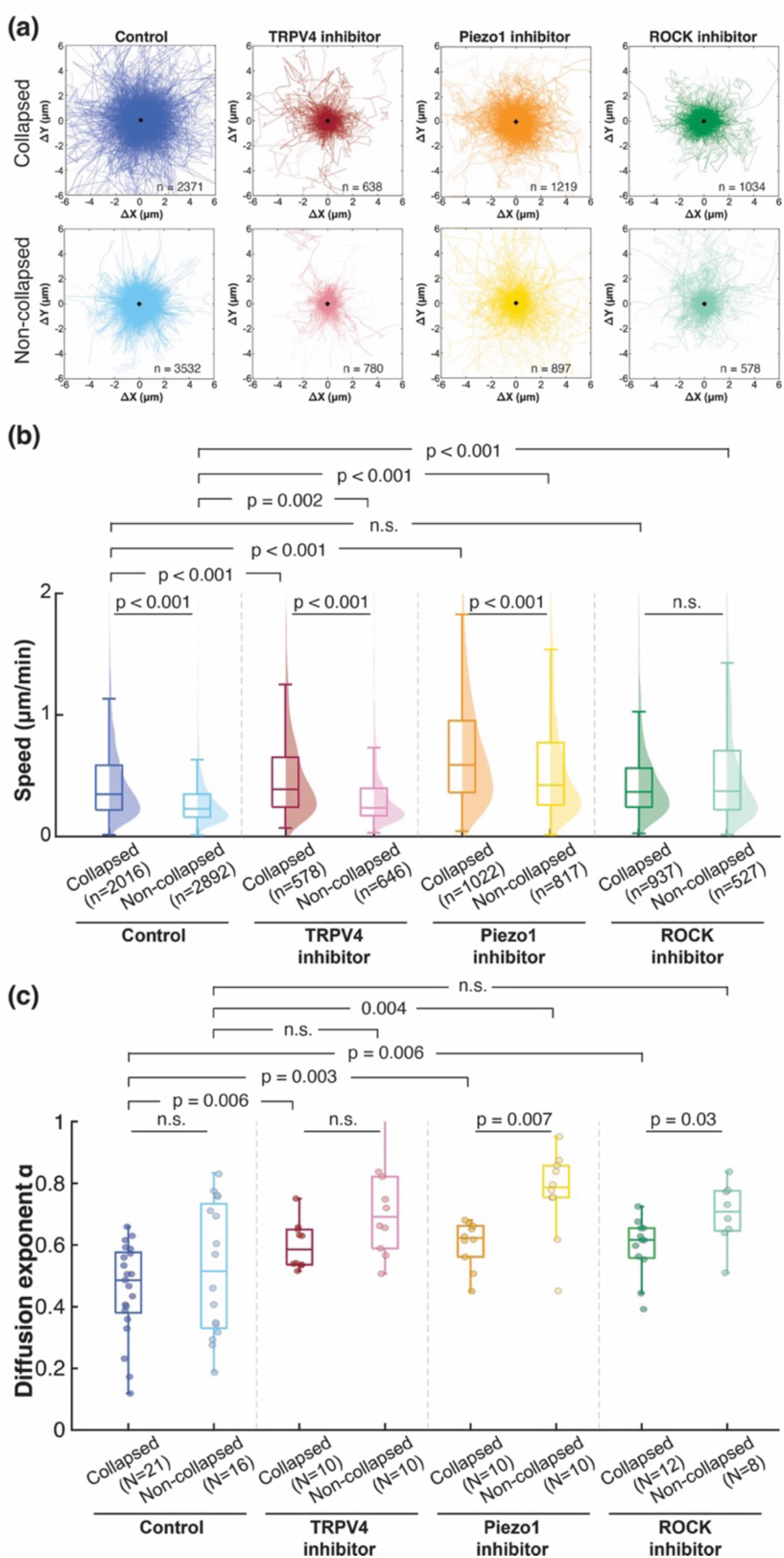
Comparison of neutrophil migration in gelatin-PCLS under the effects of migration-associated inhibitors showed the elimination of regional difference with ROCK pathway inhibition. (a) Trajectories of the last 30 min tracking (4.5-5 hours after collapsing) in all conditions. (b) Distribution of all neutrophil migration speeds. (c) Comparison of the MSD exponent α across all ROIs in gPCLS with different inhibitors. Each dot represents the α value from one ROI. Boxplot: center line, median; box edges, 25th and 75th percentiles; whiskers, 1.5× interquartile range; N: number of ROI; n: number of cell tracking. Mice = 5. For better visualization, data points exceeding the y-axis limit were excluded for plotting but included for statistical comparisons.

ROCK inhibition completely abolished the regional differences in migration speeds (Fig. 4b), with neutrophils in the collapsed and non-collapsed regions exhibiting the same median speed after treatment. ROCK is essential for actomyosin contractility by phosphorylating myosin light chain and regulating rear retraction during cell migration ^33,34^. The disappearance of regional migration differences following ROCK inhibition therefore suggests that actomyosin contractility is a critical mechanism enabling neutrophils to adapt their migration speed to the local ECM environment. This result further confirmed that the observed regional differences in gPCLS arise from neutrophil responses to tissue stiffness and/or biochemical cues, rather than from tissue geometric confinement. Interestingly, under ROCK inhibitor treatment, the migration speed in the non-collapsed region (0.37±0.03 µm/min) was significantly higher (p < 0.001), while the migration speed distribution did not change in the collapsed region (0.37±0.01 µm/min). This indicates that direct inhibition of the Rho pathway plays a smaller role in neutrophil migration in the absence of ECM stretch. Alternatively, the ROCK inhibitor can also reduce stress-fiber formation and focal adhesion maturation which may allow enhanced motility and hence higher speeds ^35^. Consistent with a contractility-dependent mechanism, inhibiting ROCK increased MSD exponent α only in the collapsed region. Since ROCK is a primary mediator of actomyosin contractility, it is likely that cell contraction primarily contributes to the subdiffusive neutrophil migration under alveolar collapse. The absence of a ROCK inhibitory effect in the non-collapsed region suggests that contractility is not the dominant factor limiting migration when the ECM is under stretch. Other mechanisms, for example cell-substrate binding, may also play a prominent role ^36^.

Inhibiting either Piezo1 or Transient receptor potential vanilloid 4 (TRPV4) channel did not eliminate the migration speed differences between collapsed and non-collapsed regions (Fig. 4b, p < 0.001). Both Piezo1 and TRPV4 are mechanosensitive calcium channels that respond to mechanical stimulation: Piezo1 is sensitive to membrane deformation while TRPV4 responds to a range of stimuli including osmotic stress and mechanical stretch ^37^. Inhibiting these channels increased neutrophil migration in both the collapsed and non-collapsed regions, indicating the ion channels play a role in the regulation of neutrophil migration. However, treating the gPCLS with either Piezo1 or TRPV4 inhibitor did not eliminate speed differences between collapsed and non-collapsed regions (Fig. 4b), suggesting that calcium signaling through these channels is not the primary determinant of collapse-sensitive migration behavior. TRPV4 inhibitors only altered the MSD exponent α in the collapsed region (Fig. 4c, p = 0.006), while the Piezo1 inhibitor influenced both regions (p < 0.005). Since an increase in α is consistent with a tendency that the cell does not stop and reverse direction, Piezo1-mediated mechanosensing reinforces neutrophil-ECM interactions.

Together, these results point to actomyosin contractility, rather than mechanosensitive calcium signaling, as a pathway governing region-dependent neutrophil migration in PCLS.

### Regional differences persist and migration speed increases under collagenase and CSE treatment

The current gel-free gPCLS model provides a high-throughput platform to investigate neutrophil migration in collapsed versus non-collapsed regions of the PCLS under the effects of different agents (Supplementary Fig. S2). Cigarette smoke is the main cause of emphysema and COPD exacerbation. During emphysema progression, an upregulated collagenase release was reported ^11^. To test whether the collapse-dependent migration phenotype is maintained under these disease-relevant stimuli, we treated gPCLS with bacterial collagenase or CSE and compared neutrophil migration speed distributions after 5 hours post-collapse (Fig. 5). In both collagenase- and CSE-treated gPCLS, neutrophils in the collapsed regions (Fig. 5b, p<0.001) still maintained significantly higher migration speeds compared to non-collapsed regions, similar to the pattern observed in untreated control gPCLS (Fig. 5b). Hence, the influence of released tension of the ECM during alveolar collapse on neutrophil migration also exists in the presence of collagenase and CSE.

**Figure 5.**
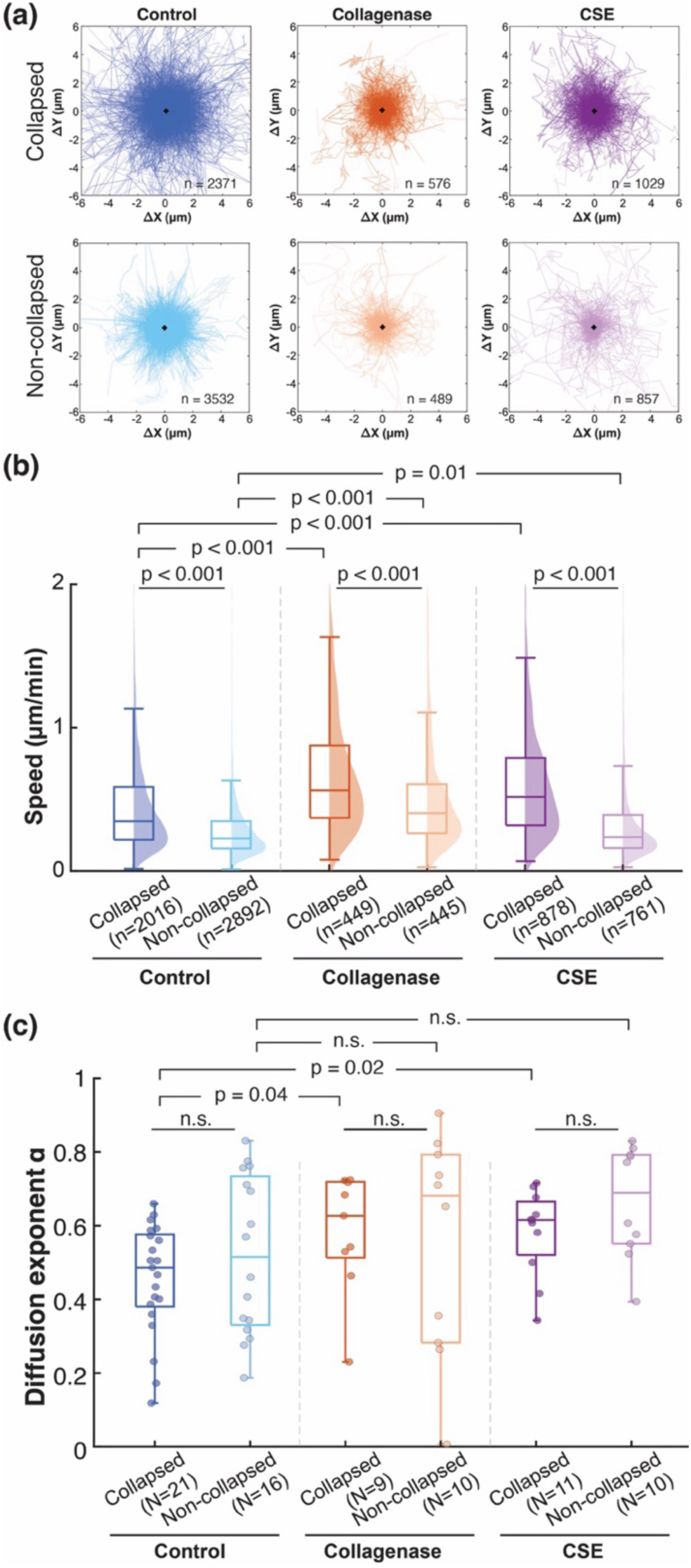
Comparison of neutrophil migration in gPCLS under effects of emphysema-associated agents revealed increased migration speed regional differences between collapsed and non-collapsed areas persisted. (a) Trajectories of the last 30 min tracking (4.5-5 hours post collapse) in all conditions. (b) Distribution of average neutrophil migration speeds within a 10-min long window at the end of cell tracking imaging. (c) Comparison of the MSD exponent α across all ROIs in gPCLS with different inhibitors. Each dot represents the α value from one ROI. Boxplot: center line, median; box edges, 25th and 75th percentiles; whiskers, 1.5× interquartile range.

Comparison of collagenase effects within each region revealed a significantly increased speed in both the collapsed and non-collapsed regions compared to the control gPCLS (Fig. 5b), suggesting that the effects of collagenase on migration is independent of mechanical stretch on lung ECM. Interestingly, collagenase also enlarged migration heterogeneity by increasing the variance of the speed distribution (p < 0.001, Fig. 5b), a different way as alveolar collapse influences speed. This speed enhancement could potentially result from two mechanisms: (1) direct enzymatic effects of collagenase on cell surface receptors ^38^, and/or (2) the generation of chemotactic collagen degradation products ^39^. The increased migration speed induced by collagenase may potentially create a positive feedback loop in which neutrophil accumulation in collapsed alveolar regions exacerbates local inflammation and drives emphysema progression through further protease release and tissue damage ^40–42^.

Similar as collagenase, CSE treatment also significantly increased migration speed in both the collapsed and the non-collapsed regions. Previous studies on isolated human neutrophils reported a decrease in migration speed following CSE exposure, due possibly to oxidative stress and impaired cytoskeletal dynamics ^43,44^. However, our results in intact gPCLS under constant CSE exposure suggest that the migration is in fact enhanced by CSE treatment. The increased migration speed under the effects of either collagenase or CSE possibly indicate an increased activation and delayed apoptosis in this inflammatory environment that can lead to greater tissue damage ^45^.

Collagenase and CSE increased MSD exponent only in collapsed region (Fig. 5c), probably by affecting the interactions between neutrophil and the ECM ^38,46^. In the non-collapsed region, neither treatment had a significant effect on the MSD exponent. Hence, we propose that in the non-collapsed region, where substrate stiffness is elevated and cell-substrate adhesion is stronger ^47^, the chemical stimuli are insufficient to overcome the enhanced binding interactions between neutrophils and the ECM. Consequently, the migratory pattern remains constrained rather than progressing toward more exploratory displacement. This reduced chemical responsiveness in the non-collapsed region may represent a protective mechanism in non-collapsed region that limits neutrophil migration and potentially restrains further inflammatory progression.

### Air-liquid interface and ventilation further enhance neutrophil migration revealed by the crystal ribcage model

To determine whether the migration behaviors observed in gPCLS translate to a more physiological setting, we employed the crystal ribcage developed by Banerji and Grifno et al. (Fig. 6a), which allows live imaging of a intact lung, observing neutrophil migration in the alveoli while preserving both the alveolar shape and the air-liquid interface under ventilation ^17^. Comparing the migration speed distribution in the collapsed region between gPCLS (liquid-filled) and crystal ribcage (air-filled) showed no difference, indicating that in the absence of alveolar stretch, the air-liquid interface alone has no effect on neutrophil migration (Fig. 6b). This suggests that the inhibitor studies found in the collapsed regions of the PCLS are applicable to those in the intact lung.

**Figure 6.**
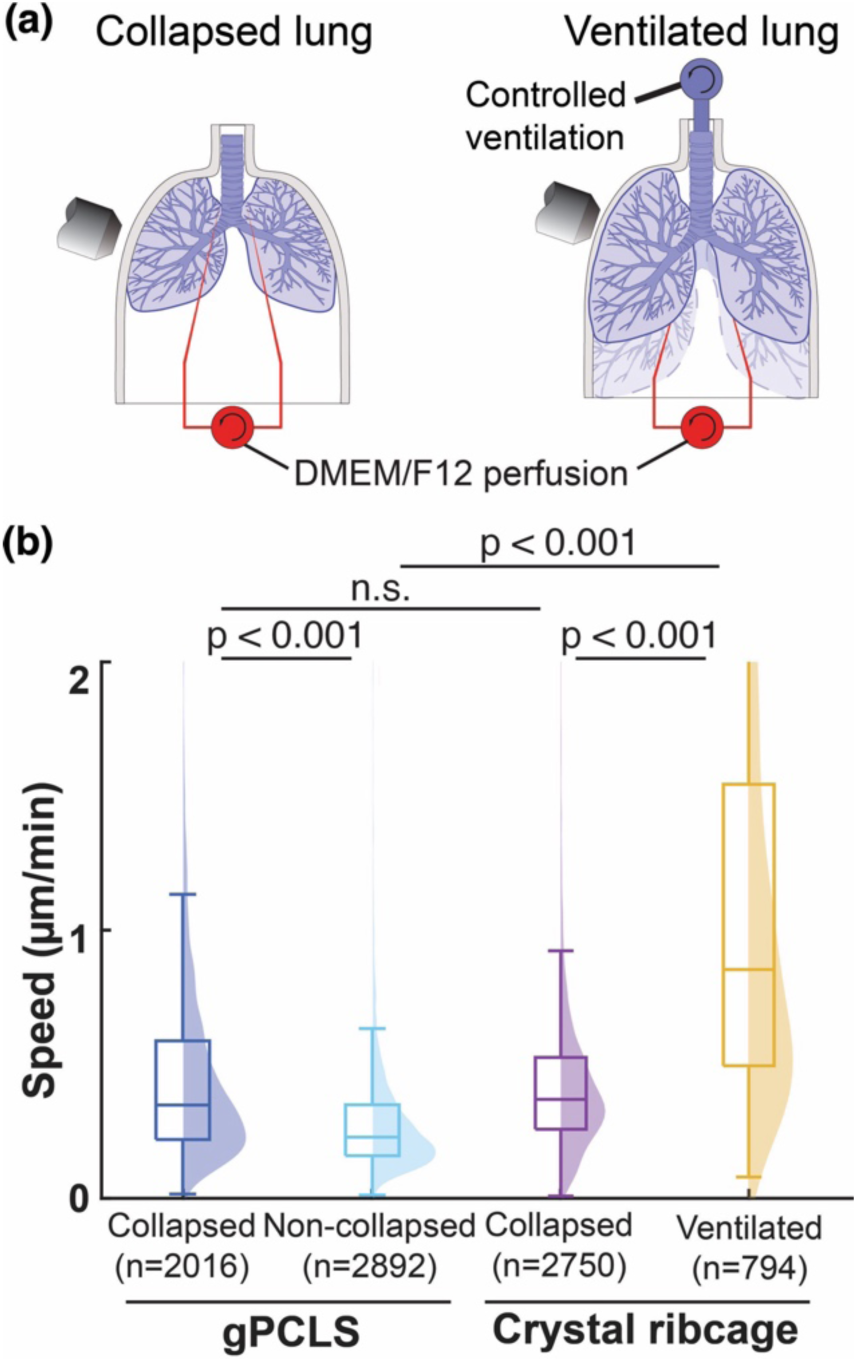
(a) For imaging in the crystal ribcage, an isolated lung without liquid filling was placed in the crystal ribcage with controlled ventilation and perfusion. Lungs were either collapsed (left) or connected to a ventilator (right), and was perfused with DMEM/F12 culture medium. (b) Comparison of neutrophil migration speeds in gPCLS and the crystal ribcage showed higher migration speeds under ventilation with preserve air-liquid interface. For better visualization, data points exceeding the y-axis limit were excluded for plotting but included for statistical comparisons.

Interestingly, with the application of dynamic ventilation superimposed on a static lung stretch in crystal ribcage, cell migration speed significantly increased to 0.85±0.05 µm/min (p < 0.001, Fig. 6b). Since surface tension of the air-liquid interface together with lung ventilation significantly affects lung stretch, we expect that mechanosensitive mechanisms likely through the Rho pathways and potentially stretch-sensitive Ca^2+^ channels underlie neutrophil migration in the lung ^8^. Khanmohammadi et al. demonstrated that cyclic stretch enhances neutrophil extracellular trap formation ^48^, which has been linked to substrate stiffness, focal adhesion, and neutrophil migration ^49^. Our findings from the crystal ribcage, where ventilation and the air-liquid interface are preserved, suggest that the combination of cyclic stretch and physiological alveolar conditions, rather than substrate stiffness alone, play a dominant role in regulating neutrophil migration in the lung parenchyma. Under static stretch in gPCLS, where these dynamic physiological conditions are absent, this enhancement of migration was not observed.

## Discussion

This study developed a gelatin-inflation method combined with a 3D-printed tissue-holding device to generate spatially defined collapsed and inflated regions within individual PCLS, enabling direct comparison of cell activities in mechanically distinct microenvironments in a high-throughput manner and under various chemical environments. PCLS can be long-term cultured and maintain functionality for up to 4 weeks ^50^ and contain all resident cells, including smooth muscle cells, epithelial cells, fibroblasts, and preserved innate and adaptive immune cells ^51^. The biological effect of long-term alveolar collapse on these resident cell populations remains unknown. For example, pulmonary dendritic cells responsible for trafficking ^52^, alveologenesis ^53^, and airway epithelial repair ^54^ have all been observed in PCLS and are known to be mechanosensitive; yet none have been investigated under heterogeneous mechanical conditions. Similarly, PCLS-based disease models of COPD ^55^ and pneumonia ^56^, as well as therapeutic development for fibrosis ^57^ and pulmonary hypertension ^58^, do not currently account for the mechanical collapse invariably present in the diseased lung. The gelatin-inflation with the tissue-holding device described here can be readily integrated with these existing PCLS applications, introducing controlled mechanical heterogeneity to reveal how alveolar collapse and stretch influence a broad range of cellular processes beyond neutrophil migration.

Several limitations should be considered when interpreting our findings. First, neutrophil phenotype and behavior differ between LPS-induced acute inflammation and chronic emphysema. While we used LPS to recruit neutrophils to the lung and create a controlled inflammatory environment, neutrophils in the emphysematous lungs exhibit altered activation states and distinct protease expression profiles compared to acutely recruited neutrophils ^1,59,60^. Second, the PCLS culture system does not permit media exchange after drug treatments, resulting in continuous and high concentration exposure to collagenase, CSE, or pharmacological inhibitors throughout the observation period. This differs from in vivo scenarios where local concentrations of agents are much smaller and fluctuate over time and space due to clearance, metabolism, and diffusion. Third, in the gelatin-filled model, the collapsed regions were consistently located near the pleural areas, while the stretched regions were close to the central locations. Regional differences in baseline immune cell distribution between peripheral and central lung regions may contribute to the observed migration patterns independent of mechanical factors. This baseline difference was observed in aPCLS where both central and pleural region were inflated with agarose. Fourth, unlike the emphysematous lung, alveolar collapse in the current model was introduced mechanically, without the complex biochemical and structural alterations in the ECM such as mechanical rupture ^61^ that characterize emphysema pathology.

Overall, our findings reveal that alveolar collapse enhances neutrophil motility, and this collapse-dependent behavior requires actomyosin contractility through the ROCK pathway. The application of disease-relevant stimuli including collagenase and CSE treatment increased the migration speed, but did not reverse the regional differences between collapsed and non-collapsed regions. Furthermore, neutrophil migration speed was accelerated by the presence of both air-liquid interface and dynamic breathing-like alveolar stretch in the crystal ribcage.

Our findings carry important implications for understanding lung disease pathogenesis. In collapsed regions of the parenchyma, neutrophil motility and displacement were consistently higher than in non-collapsed regions, a difference that persisted even under inflammatory biochemical stimuli, suggesting amplified local neutrophil-mediated tissue damage through increased proteinase release and oxidative burden. In the context of emphysema, this increased neutrophil migration in collapsed regions may reflect an upregulation of neutrophil collagenase release, which could drive further tissue destruction with subsequent airspace enlargement and alveolar collapse. This process potentially creates a positive feedback loop in which mechanical collapse promotes neutrophil activity that in turn accelerates structural remodeling. The identification of the ROCK pathway as a critical mediator of this collapse-dependent migration points to actomyosin contractility as a viable therapeutic target to interrupt this cycle and limit neutrophil-driven inflammation in the lung.

## Acknowledgments

We thank the Boston University Neurophotonics Center and Boston University Micro Nano Imaging Core Facility for their generous support and access to their imaging resources, particularly the Nikon W1 SoRa spinning disk microscope acquired through NSF MRI Award #2215990. We acknowledge the following research support: NIH R21EB031332 and DP2HL168562, Sloan Research Fellowship, Beckman Young Investigator Award, NSF CAREER, Boston University CMTM and Dean’s Catalyst Awards, and the American Cancer Society Institutional Fund at Boston University to H.T.N; Kilachand Fund to H.T.N. and J.P.M.; NIH F30HL168952 to L.S.; NSF GRFP to G.N.G. and L.C.; NIH S10OD024993 to Boston University BME Department. National Science Foundation (NSF) CAREER Award (CMMI-2047683; J.W.B.), and the AFOSR DURIP award (FA9550-24-1-0073).

