## Supplementary for "Neutrophil migration in the lung is altered by alveolar collapse and stretch"

* Corresponding authors.

Supplemental methods

*Animal use ethics*

Male mT/mG mice (B6.129(Cg)-Gt(ROSA)26Sortm4(ACTB-tdTomato,-EGFP)Luo/J; JAX stock #007676, Jackson Laboratory) aged 16–20 weeks were used for all experiments. A breeding pair was purchased from Jackson Laboratory to establish a colony, which served as the source for all experimental animals.

All experiments conformed to ethical principles and guidelines under protocols approved by the Boston University Institutional Animal Care and Use Committee. Mice were housed and bred under pathogen-free, ambient temperature, humidity and light–dark conditions at the Boston University Animal Science Center. No housing or handling exceptions were made for this study.

*Preparation of CSE*

CSE solutions were prepared and used following a previously established protocol ^1^. Briefly, two cigarettes (Marlboro Red, Philip Morris USA, Richmond, VA, United States) with filters removed were bubbled through 20 mL of DMEM/F12 at a flow rate of 2.5 L/min to yield a stock solution of 0.1 cig/mL. The solution was immediately sterile-filtered using a 0.22 μm pore size syringe filter to remove tobacco debris and particulates. Stock solutions were aliquoted and frozen at ‑20°C until use. For experiments, the stock solution was diluted in culture medium to achieve a working concentration of 0.008 cig/mL.

*Reagent preparation*

Stock solutions of reagents were prepared and stored as described in Supplementary Table S1.

*Mouse model*

To recruit neutrophils to the lung, mice (body weight ~30 g) received an intraperitoneal injection of LPS (Supplementary Table S1) 7 hours prior to euthanasia ^2^. To enable fluorescence imaging of neutrophils in PCLS, Ly-6G antibody (Supplementary Table S1) was administered via retro-orbital injection 3.5 hours prior to euthanasia. This timing allowed for neutrophil recruitment and labeling prior to PCLS generation (Supplementary Fig. S2).

*Lobar injection of agarose and gelatin solution into the same lung*

All surgical instruments were sterilized by autoclaving. Non-autoclavable components (e.g., catheter tubing) were sterilized by UV exposure for at least 20 minutes prior to use. A 6% (w/v) gelatin solution (Sigma-Aldrich, G2500) and 1.5% (w/v) low-melting point agarose (Sigma-Aldrich, A2576) were maintained at 37°C in a water bath before inflation. The experimental timeline is summarized in Supplementary Fig. S2. Briefly, heparin was injected intraperitoneally 15 min before euthanasia was performed via CO_2_ asphyxiation or via isoflurane overdose followed by cervical dislocation. The chest cavity and rib cage were then opened above the diaphragm, allowing the lung to collapse spontaneously. A small incision was made in the upper portion of the trachea to create an opening for cannulation. Two 24G x 0.75in catheters were connected via catheter tubing to two 10 mL syringes filled with 37°C agarose or gelatin solution (Fig. 1a). Each cannula was then inserted from the proximal tracheal end angled into the left or right mainstem bronchus and ligated with sutures to seal off the bronchi around the catheters, respectively.

The lung was inflated using two syringe pumps set to deliver agarose or gelatin solutions at a controlled rate of 0.02 mL/s. This slow inflation rate allowed sufficient time for gelatin to diffuse throughout the airways and achieve uniform lung inflation. Inflation was stopped based on visual evaluation of the size of the lung. Immediately following inflation, the surface of the lung was flushed with chilled Hank's Balanced Salt Solution (HBSS) to rapidly solidify the agarose/gelatin within the alveolar spaces. The lung-heart block was then excised and placed in chilled HBSS with 20mM HEPES. One agarose-inflated lobe and one gelatin-inflated lobe were carefully dissected and used for PCLS preparation. The lobe assigned to each gel inflation type inflation was alternated across experiments.

*PCLS generation*

aPCLS and gPCLS were generated separately from the lobes filled with agarose and gelatin, respectively. 250 μm-thick PCLS were generated using Compresstome VF-300-0Z vibrating microtome (Precisionary Instruments, St. Louis, MO, United States) with the lobe embedded in 1.5% agarose on the specimen holder. The sectioning was done at 4°C using the highest oscillation frequency and lowest advancement speed settings to minimize tissue deformation. PCLS were collected in chilled HBSS supplemented with 20 mM HEPES buffer and maintained at 4°C before use.

*Tissue-holding device printing*

The tissue-holding device was designed using SolidWorks (Dassault Systèmes) and fabricated using a high-resolution 3D printer (microArch S240, Boston Micro Fabrication) and HTL resin (Boston Micro Fabrication) at a printing resolution of 20 µm in z and 10 µm in x and y (Fig. 1b, i). The exposure time and power intensity level were 1.0 s and 50 (out of 255), respectively. The support structures were generated using VoxelDance Additive software (VoxelDance) based on the parameters provided in the manufacturer's manual for the microArch S240 and HTL resin. Each tissue-holding device consisted of 24 needles arranged on the inner edge of an elliptical base (2.5 × 5 mm, 1 mm wall width) with a base thickness of 1 mm. Individual needles were designed with a tip diameter of 0.05 mm, a base diameter of 0.3 mm, and a length of 0.5 mm (Fig. 1b, i and ii). The printed devices were washed in 70% isopropyl alcohol (Sigma-Aldrich, 190764) to remove the residual resin and post-cured using a Formcure (Formlabs) at 60°C for 3 hours.

*PCLS mounting and treatment*

Immediately after PCLS were generated from both lobes, individual PCLSs were transferred to a 12-well imaging plate and positioned with custom 3D-printed tissue-holding device (Fig. 1b) that mechanically supported a region of the slice. A stainless steel M2 nut was placed on top of each tissue-holding device to provide additional weight and prevent sample drifting. The pin-PCLS assembly was further embedded in a 37°C liquid solution of 1:1 mixture of 3% agarose and DMEM/F12 culture medium. The agarose/medium mixture was rapidly solidified by placing the plate on ice to stabilize the sample during long-term live imaging. After solidification, 500 μL of DMEM/F12 culture medium was added to each well.

CSE or bacterial collagenase (Supplementary Table S1) were added to the culture medium of gPCLS prior to incubation (Supplementary Table S1). gPCLS were then incubated at 37°C with 5% CO₂ for 30 minutes to allow gelatin melting, which resulted in alveolar deflation and collapse outside the pin-supported region while inside the pin-secured region tissue remained mechanically stretched (Fig. 1d, i). For experiments involving inhibitors (TRPV4, ROCK, or Piezo1 inhibitors), gPCLS were first imaged for 1 hour to establish baseline neutrophil migration, after which inhibitors were added to the medium at their respective concentrations (Supplementary Table S1).

*Live imaging of neutrophil migration*

Neutrophil migration in PCLS was imaged using a confocal microscope (Nikon CSU-W1 SoRa spinning disk confocal microscope, Nikon Instruments, Melville, NY, United States) equipped with a stage-top incubation chamber maintained at 37°C with 5% CO₂.

For each PCLS, four regions of interest (ROIs) were selected: 2 outside the pin-supported area (collapsed region in gPCLS) and 2 inside the pin-supported area (non-collapsed region in gPCLS). Timelapse imaging was performed by sequentially acquiring all ROIs across all PCLS in a repeating cycle, yielding a temporal resolution of 1 minute per frame for each individual ROI (Supplementary Fig. S2). Imaging was conducted using a 10x objective with dual-channel fluorescence acquisition: 640 nm laser excitation to visualize Alexa Fluor® 647-labeled neutrophils (Ly-6G+) and 516 nm laser excitation to visualize the mT/mG membrane signal for alveolar structure. Single focal plane images were acquired at each time point for each ROI. The total imaging time was up to 5 hours.

*Image stabilization*

To correct sample drift during time-lapse imaging due to thermal fluctuations and mechanical perturbations, a two-step image stabilization workflow was implemented using custom MATLAB (The MathWorks, Natick, MA, United States) and Python scripts.

The mTmG membrane signal from the lung alveolar structure was used as a reference to calculate frame-to-frame motion. First, global motion correction was performed using a similarity transformation algorithm in MATLAB (estimateGeometricTransform2D function) to align the structural channel across all frames, producing a relatively stable video with large-scale drifts removed. Second, residual local motion was corrected using an optical flow-based algorithm implemented in Python with the OpenCV library (calcOpticalFlowFarneback function), which performed fine-scale, localized stabilization to account for non-rigid tissue deformations.

The motion correction transformations calculated from the structural channel (mT/mG signal) were applied identically to the neutrophil channel at each step to maintain spatial correspondence between channels. This ensured that neutrophil positions were accurately tracked relative to the underlying tissue architecture throughout the imaging period.

*Neutrophil migration quantification*

Neutrophil detection and tracking were performed using Imaris software (v9.8.2, Oxford Instruments, Bitplane, Zürich, Switzerland). Individual neutrophils were identified using the Spots detection algorithm (Fig. 2a). Regions exhibiting image distortion artifacts from the stabilization process were manually excluded from the analysis to ensure accurate tracking. Neutrophil trajectories were reconstructed across time-lapse sequences using the autoregressive motion tracking algorithm. The software generated quantitative output statistics for each tracked neutrophil, including instantaneous migration speed, mean migration speed, track displacement, and cell position coordinates at each time point. Tracks shorter than 3 minutes were excluded from analysis to eliminate tracking artifacts.

For each experimental condition, neutrophil instantaneous migration speeds were plotted in a speed-time plot, with each curve smoothed over a 10-min window. Statistical analysis of the migration speeds was performed at the end of the imaging period, where cell motility reached a stable regime and differences between conditions were most pronounced were to compare differences between groups. For speed distribution plots, an upper threshold was applied for better visualization while the speed values were included in statistical comparisons.

MSD, a measure of migration type, was calculated based on the cell tracking trajectory (Fig. 2d, i). For each region of interest (ROI), the MSD as a function of time lag $\Delta t$ was computed as

$MSD=\langle[\vec{r}(t + \Delta t) - \vec{r}(t)]^{2}\rangle$*,*

where $\vec{r}$ is the position of the neutrophil at time t, and <…> brackets represent time averaging over all valid displacement pairs separated by $\Delta t$ across all tracked neutrophils within the ROI. Displacements were identified by matching time points within the time lag window on each $\Delta t$ (Fig. 2d, i). In anomalous diffusion, the MSD is described as

$MSD \propto\Delta t^{\alpha}$*,*

where $\alpha$ is the MSD exponent calculated as the slope of a linear fit on a log-log plot of MSD versus time lag. The fitting was performed over time lags of 1 to 30 min (Fig. 2d, ii), as extending beyond this range resulted in a biased analysis on longer tracks. A value of $\alpha$ = 1 corresponds to Brownian motion whereas $\alpha$ < 1 indicates a subdiffusive process where internal or external factors restrict movement.

*Random walk simulation of neutrophil migration in PCLS*

Tissue images were loaded into MATLAB (2024a) and resized to 60% of the original scale to limit computational burden. Contrast within images was then increased by using MATLAB’s histeq function. Images were then automatically binarized using MATLAB’s imbinarize function to capture the tissue geometry, in which white pixels corresponded to walkable tissue space. Only the largest contiguous structure in the image was kept for the random walks.

Next, 1000 Monte Carlo random walkers (about 10 times bigger than experimental data size) were simulated on each binary image (Supplementary Fig. S3), representing neutrophil migration in PCLS. Each walk started on a random tissue pixel and was allowed to move to any connected tissue pixel (up, down, left, right). 500 steps were allowed for each walker. The random walk MSD and exponent α was computed using the same method applied to experimentally tracked neutrophils.

*Trajectory preprocessing*

To ensure analytical consistency and mitigate variability arising from differences in trajectory duration and orientation, all cell trajectories were standardized to a fixed temporal window of 30 minutes (Fig. 2, green box). Trajectories containing missing time points were excluded and only cells continuously tracked over the entire interval were retained for downstream analysis to ensure data robustness. To account for directional heterogeneity, we established an intrinsic migration coordinate framework. The primary migration axis was defined as the direction of maximal cumulative displacement, with a secondary axis defined orthogonally. These axes were derived using singular value decomposition (SVD) of the displacement matrix $A$, such that:

$$A=U\Sigma V^{T}$$

where $U$ and $V^{T}$ contain the eigenvectors of $AA^{T}$ and $A^{T}A$, respectively, and $\Sigma$ is a diagonal matrix of singular values. The original coordinate matrix $R$ was then projected onto this orthonormal basis via $V$, such that:

$$R_{\text{reg}}=RV$$

Within this transformed coordinate system, the first and second columns of $R_{\text{reg}}$ correspond to the trajectories along the primary and secondary migration axes, respectively.

*Multidimensional motility analysis.*

To quantitatively characterize single-cell motility behaviors, we computed a set of 93 features capturing multiple dimensions of movement dynamics across 7,938 neutrophil tracking. These features encompass both displacement- and turning angle–based metrics, reflecting properties such as movement magnitude, spatial and temporal organization, signal characteristics, correlation patterns, entropy, and decomposed motion components ^3,4^. A comprehensive list of motility features is provided in Supplementary Table S2. Then, principal component analysis (PCA) was applied to reduce redundancy among correlated variables. Principal components (PCs) accounting for 95% of the total variance were retained for subsequent analysis. These PCs were then used as inputs for two-dimensional embedding via UMAP, followed by unsupervised clustering. Clustering was performed using the K-means++ algorithm, and cluster validity was assessed by visualizing trajectory projections associated with each MC to evaluate qualitative consistency. To ensure statistical robustness, clusters containing fewer than 100 cells were excluded as outliers, resulting in eight distinct MCs. Furthermore, Shannon entropy (S) was computed to quantify the degree of heterogeneity across motility states, as defined below:

$$S=\sum_{i=1}^{8} -p_{i}\cdot\log\left( p_{i} \right)$$

Here $p_{i}$ is the fraction of cells within each MC $i$.

*Software*

Multidimensional motility analysis was performed in Python with the following software specifications: python 3.9.13, scipy 1.13.1, scikit-learn 1.1.13, scikit-image 0.19.3, scikit-posthocs 0.9.0, statsmodels 0.13.5, pandas 1.5.2, matplotlib 3.6.2, seaborn 0.11.2, umap-learn 0.5.3, and numpy 1.23.5, cmcrameri 1.9, EntropyHub 0.2, pyarrow 12.0.1.

*Crystal ribcage design and fabrication*

The crystal ribcage ^5^ provides a transparent, biocompatible physiological environment that is matched to the age and strain of the animal for ex vivo lung imaging. Briefly, a three-dimensional model of the thoracic cavity was generated using micro-computed tomography (µCT) scans of the C57BL/6 mouse chest. This model was then printed using a stereolithography 3D printer (Form3, FormLabs). After printing, the surface was polished to remove printing artifacts and the printed model was used as a positive mold to thermoform the crystal ribcage.

To provide a stable low-friction interface that minimizes mechanical resistance during lung ventilation, the inner surface of the crystal ribcage was dip-coated with iSurGlide Plus (iSurTech) which provides a lubricated and optically transparent hydrophilic surface ^6^.

*Imaging of neutrophil in ex vivo lung inside crystal ribcage*

The timeline for neutrophil imaging in the crystal ribcage is described in Supplementary Figure S2. Briefly, mice that had been treated with LPS as described above were anesthetized with a ketamine/xylazine cocktail (100 and 10 mg kg⁻¹, respectively). After induction of anesthesia, the abdominal cavity was exposed and 100 µL of 2.5 mg/mL heparin was administered intrahepatically (Supplementary Fig. S2).

A tracheal cannula was inserted to the lung and connected to a ventilator (RoVent, Kent Scientific) with ventilation rate at 120 breaths per minute with airway pressures between 5 and 10 cmH₂O. The mouse was then euthanized by exsanguination following transection of the caudal vena cava.

The lungs were perfused with DMEM/F12 medium at 20 µL/min through two cannulas inserted into the pulmonary artery and left atrium. After at least 10 minutes of perfusion under ventilation, the lung–heart block was removed from the thoracic cavity and placed into the crystal ribcage for microscopy (Fig. 6a).

Neutrophils in the ex vivo lung inside the crystal ribcage were imaged under collapsed or ventilated conditions using an upright spinning-disk confocal microscope (Nikon Crest X-Light V3) with a 10× objective at 37 °C (Fig. 6a). Imaging was taken at 30-second time intervals to improve image stabilization in post-processing. For imaging in the collapsed condition, the lung was collapsed before placing into the crystal ribcage. For imaging in the non-collapsed condition, the lung was ventilated continuously with a pause for 5s at a pressure of 14 cmH₂O in each image acquisition. In both cases, the lung was perfused with DMEM/F12 culture medium at a flow rate of 20 µL/min. The imaging frames were extracted at every 2nd frame to match the frame rate of 1 min per frame in PCLS imaging.

Figures

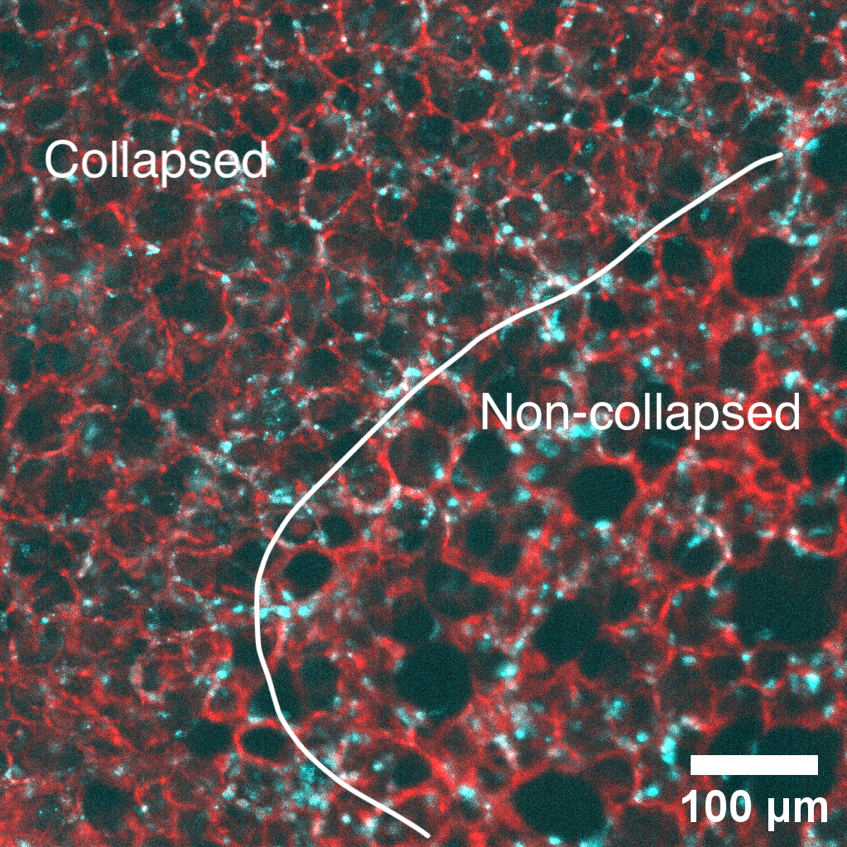

**Supplementary figure S1.** Inhomogeneous inflation where agarose failed to diffuse deeper into the distal side of the airway (left), generating collapsed and non-collapsed region within the same aPCLS. White line: boundary between the 2 regions.

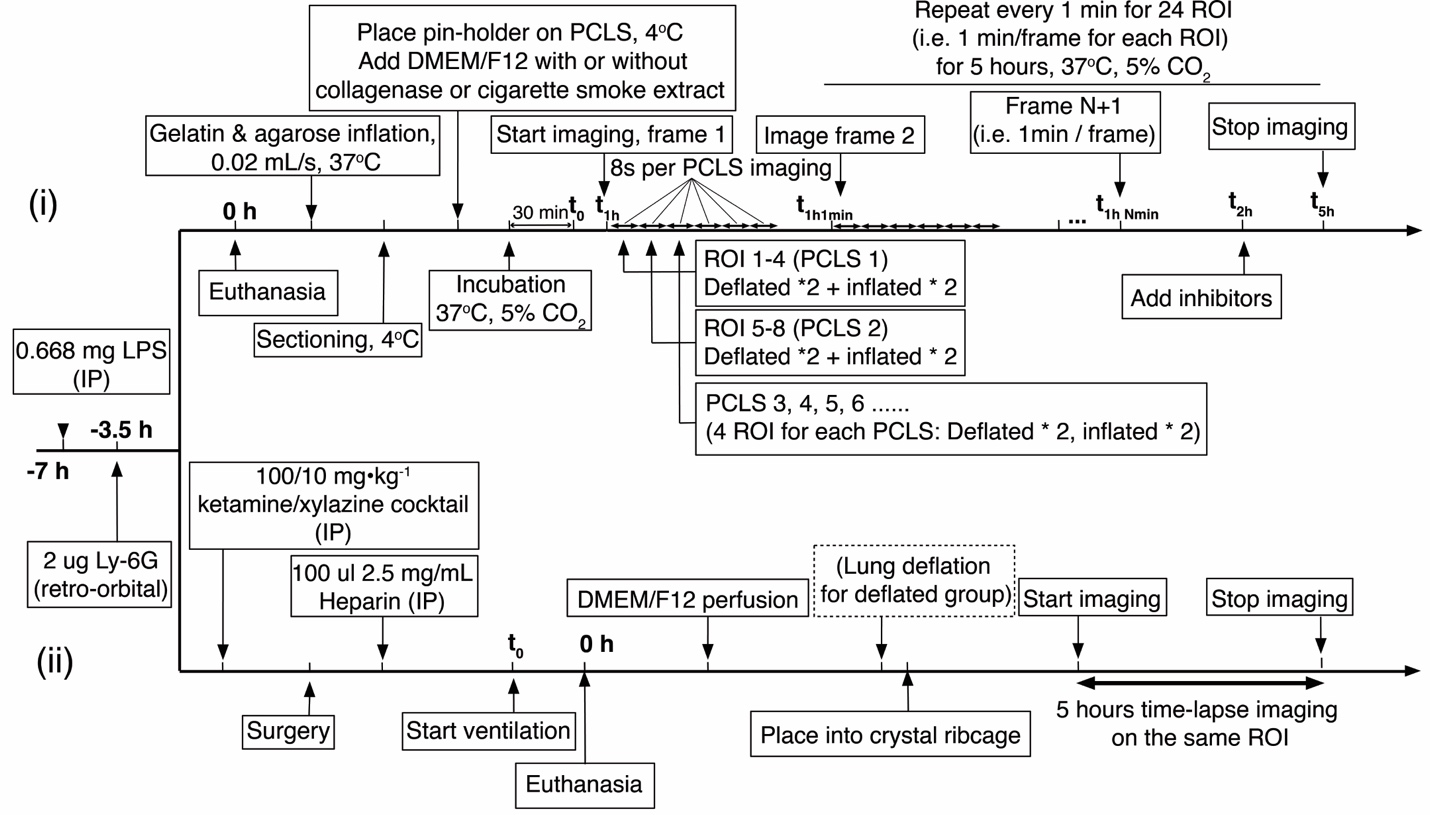

**Supplementary figure S2.** Experiment timeline. (i) PCLS imaging protocol. (ii) crystal ribcage imaging protocol. t0 in each timeline marks the onset of post-collapse time (i.e., the origin of the time axis in all speed-versus-time analyses). IP: intraperitoneal

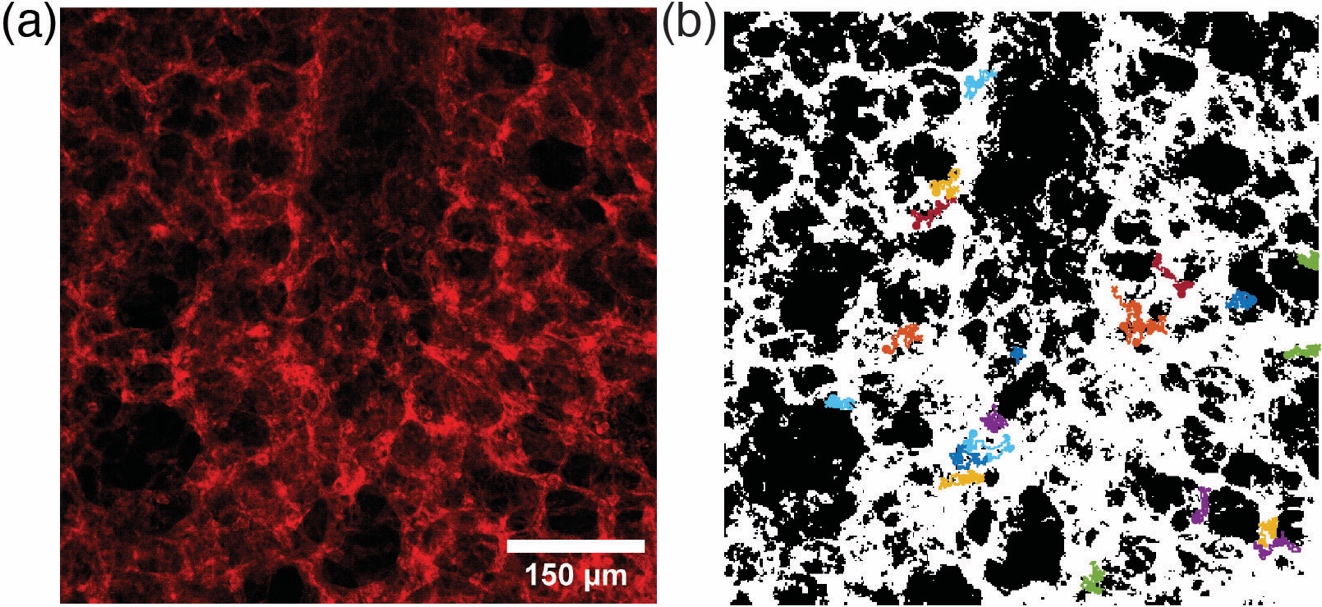

**Supplementary Figure S3.** (a) PCLS tissue geometry and (b) simulated neutrophil migration on binarized image.

Supplementary Table S1 Reagent concentration and preparation

| Reagent | Catalog number | Solvent | Stock  conc. | Working conc. /Dose |
| --- | --- | --- | --- | --- |
| Lipopolysaccharide from E. coli O55:B5 | Sigma-Aldrich, L2880 | PBS | 3.3 mg/mL | 0.67 mg ^5^ |
| Alexa Fluor® 647 anti-mouse Ly-6G antibody | BioLegend, 127610 | PBS | 40 μg/mL | 2 μg |
| Piezo1 inhibitor GsMTx4 | MedChemExpress, HY-P1410 | PBS | 200 μM | 8 μM ^7^ |
| TRPV4 inhibitor GSK2193874 | MedChemExpress, HY-100720 | DMSO | 2.3 mM | 4.6 μM^8^ |
| ROCK inhibitor Y-27632 | MedChemExpress, HY-10583 | PBS | 2.5 mM | 5 μM ^9^ |
| Bacterial collagenase from Clostridium histolyticum | Sigma-Aldrich, C0130 | PBS | 10 mg/mL | 0.4 mg/mL ^10^ |

**Supplementary Table S2.** Motility features.

| Index | Parameter | Type | Description |
| --- | --- | --- | --- |
| 1 | total_distance | Displacement | Sum of all stepwise displacements (path length). |
| 2 | avg_speed | Displacement | Mean speed over the trajectory. |
| 3 | max_speed | Displacement | Maximum instantaneous speed. |
| 4 | min_speed | Displacement | Minimum instantaneous speed. |
| 5 | net_distance | Displacement | Straight line distance from start to end. |
| 6 | progressivity | Turning angle | Path straightness (net distance divided by total distance). |
| 7 | total_angle | Turning angle | Total turning angles accumulated across steps. |
| 8 | avg_angle | Turning angle | Mean turning angle between successive steps. |
| 9 | max_angle | Turning angle | Maximum instantaneous turning angle. |
| 10 | min_angle | Turning angle | Minimum instantaneous turning angle. |
| 11 | displ_variance | Distribution descriptor | Variance of the displacement distribution from the trajectory |
| 12 | displ_cov | Distribution descriptor | Coefficient of variation of displacement (standard deviation divided by the mean). |
| 13 | displ_skewness | Distribution descriptor | Skewness of the displacement distribution from the trajectory. |
| 14 | displ_kurtosis | Distribution descriptor | Kurtosis of the displacement distribution from the trajectory. |
| 15 | displ_ngaussalpha | Distribution descriptor | Non Gaussian parameter for the displacement distribution from the trajectory. |
| 16 | displ_gini | Distribution descriptor | Gini coefficient of inequality in displacements. |
| 17 | angle_variance | Distribution descriptor | Variance of the turning angle distribution from the trajectory |
| 18 | angle_cov | Distribution descriptor | Coefficient of variation of turning angle (standard deviation divided by the mean). |
| 19 | angle_skewness | Distribution descriptor | Skewness of the turning angle distribution from the trajectory. |
| 20 | angle_kurtosis | Distribution descriptor | Kurtosis of the turning angle distribution from the trajectory. |
| 21 | angle_ngaussalpha | Distribution descriptor | Non Gaussian parameter for the turning angle distribution from the trajectory. |
| 22 | angle_gini | Distribution descriptor | Gini coefficient of inequality in turning angles. |
| 23 | alphas | Mode | Anomalous diffusion exponent from MSD scaling. |
| 24 | msds_1 | Displacement | Mean squared displacement at time lag = 1. |
| 25 | msds_2 | Displacement | Mean squared displacement at time lag = 2. |
| 26 | msds_3 | Displacement | Mean squared displacement at time lag = 3. |
| 27 | displ_autocorr_1 | Temporal correlation | Autocorrelation of displacements at time lag = 1. |
| 28 | displ_autocorr_2 | Temporal correlation | Autocorrelation of displacements at time lag = 2. |
| 29 | displ_autocorr_3 | Temporal correlation | Autocorrelation of displacements at time lag = 3. |
| 30 | displ_partial_autocorr_1 | Temporal correlation | Partial autocorrelation of displacements at time lag = 1. |
| 31 | displ_partial_autocorr_2 | Temporal correlation | Partial autocorrelation of displacements at time lag = 2. |
| 32 | displ_partial_autocorr_3 | Temporal correlation | Partial autocorrelation of displacements at time lag = 3. |
| 33 | angle_autocorr_1 | Temporal correlation | Autocorrelation of turning angles at time lag = 1. |
| 34 | angle_autocorr_2 | Temporal correlation | Autocorrelation of turning angles at time lag = 2. |
| 35 | angle_autocorr_3 | Temporal correlation | Autocorrelation of turning angles at time lag = 3. |
| 36 | angle_partial_autocorr_2 | Temporal correlation | Partial autocorrelation of turning angles at lag 2. |
| 37 | angle_partial_autocorr_3 | Temporal correlation | Partial autocorrelation of turning angles at lag 3. |
| 38 | avg_speed_c | Decomposed features | Mean speed projected onto the primary migration axis. |
| 39 | max_speed_c | Decomposed features | Maximum instantaneous speed along the primary axis. |
| 40 | min_speed_c | Decomposed features | Minimum instantaneous speed along the primary axis. |
| 41 | net_distance_c | Decomposed features | Net displacement along the primary migration axis. |
| 42 | progressivity_c | Decomposed features | Straightness along the primary axis: net primary displacement divided by total primary path length. |
| 43 | avg_speed_l | Decomposed features | Mean instantaneous speed projected onto the secondary axis. |
| 44 | max_speed_l | Decomposed features | Maximum instantaneous speed along the secondary axis. |
| 45 | min_speed_l | Decomposed features | Minimum instantaneous speed along the secondary axis. |
| 46 | net_distance_l | Decomposed features | Net displacement along the seoncdary axis. |
| 47 | progressivity_l | Decomposed features | Straightness along the secondary axis: net secondary displacement divided by total secondary path length. |
| 48 | Max_distance_l / Max_distance_c | Decomposed features | Ratio of maximum secondary distance to maximum primary distance. |
| 49 | Distance_l / Distance_c | Decomposed features | Ratio of total seoncdary path length to total primary path length. |
| 50 | msd_c_1 | Decomposed features | Mean squared displacement of the primary-axis component at time lag = 1. |
| 51 | msd_c_2 | Decomposed features | Mean squared displacement of the primary-axis component at time lag = 2. |
| 52 | msd_c_3 | Decomposed features | Mean squared displacement of the primary-axis component at time lag = 3. |
| 53 | msd_l_1 | Decomposed features | Mean squared displacement of the secondary-axis component at time lag = 1. |
| 54 | msd_l_2 | Decomposed features | Mean squared displacement of the secondary-axis component at time lag = 2. |
| 55 | msd_l_3 | Decomposed features | Mean squared displacement of the secondary-axis component at time lag = 3. |
| 56 | displ_variance_c | Decomposed features | Variance of the displacement distribution from the primary axis |
| 57 | displ_cov_c | Decomposed features | Coefficient of variation of displacement from the primary axis (standard deviation divided by the mean). |
| 58 | displ_skewness_c | Decomposed features | Skewness of the displacement distribution from the primary axis. |
| 59 | displ_kurtosis_c | Decomposed features | Kurtosis of the displacement distribution from the primary axis. |
| 60 | displ_ngaussalpha_c | Decomposed features | Non Gaussian parameter for the displacement distribution from the primary axis. |
| 61 | displ_variance_l | Decomposed features | Variance of the displacement distribution from the secondary axis |
| 62 | displ_cov_l | Decomposed features | Coefficient of variation of displacement from the secondary axis (standard deviation divided by the mean). |
| 63 | displ_skewness_l | Decomposed features | Skewness of the displacement distribution from the secondary axis. |
| 64 | displ_kurtosis_l | Decomposed features | Kurtosis of the displacement distribution from the secondary axis. |
| 65 | displ_ngaussalpha_l | Decomposed features | Non Gaussian parameter for the displacement distribution from the secondary axis. |
| 66 | displ_autocorr_c_1 | Decomposed features | Autocorrelation of displacements along the primary axis at time lag = 1. |
| 67 | displ_autocorr_c_2 | Decomposed features | Autocorrelation of displacements along the primary axis at time lag = 2. |
| 68 | displ_autocorr_c_3 | Decomposed features | Autocorrelation of displacements along the primary axis at time lag = 3. |
| 69 | displ_autocorr_l_1 | Decomposed features | Autocorrelation of displacements along the secondary axis at time lag = 1. |
| 70 | displ_autocorr_l_2 | Decomposed features | Autocorrelation of displacements along the secondary axis at time lag = 2. |
| 71 | displ_autocorr_l_3 | Decomposed features | Autocorrelation of displacements along the secondary axis at time lag = 3. |
| 72 | inst_speed_peak_to_peak | Signal descriptors | Peak-to-peak amplitude of the instantaneous speed series. |
| 73 | inst_speed_slopes | Signal descriptors | Mean absolute slope (discrete derivative) of the instantaneous speed series. |
| 74 | inst_speed_RMSs | Signal descriptors | Root-mean-square (RMS) of the instantaneous speed series. |
| 75 | inst_speed_crestfactor | Signal descriptors | Crest factor of instantaneous speed (peak divided by RMS). |
| 76 | inst_speed_formfactor | Signal descriptors | Form factor of instantaneous speed (RMS divided by mean absolute value). |
| 77 | inst_speed_pulseindicator | Signal descriptors | Pulse indicator of instantaneous speed (peak divided by mean absolute value). |
| 78 | inst_speed_approximate_entropies | Entropy | Approximate entropy of the instantaneous speed series (pattern regularity). |
| 79 | inst_speed_attention_entropies | Entropy | Entropy of attention-weight distributions derived from the instantaneous speed series. |
| 80 | inst_speed_permutation_entropies | Entropy | Permutation entropy of the instantaneous speed series. |
| 81 | inst_speed_bubble_entropies | Entropy | Bubble entropy, a noise-robust complexity metric of the instantaneous speed series. |
| 82 | inst_speed_cosine_similarity_entropies | Entropy | Entropy of cosine-similarity distributions between windowed segments of the speed series. |
| 83 | inst_angle_peak_to_peak | Signal descriptors | Peak-to-peak amplitude of the instantaneous turning-angle series. |
| 84 | inst_angle_slopes | Signal descriptors | Mean absolute slope of the instantaneous turning-angle series. |
| 85 | inst_angle_RMSs | Signal descriptors | Root-mean-square (RMS) of the instantaneous turning-angle series. |
| 86 | inst_angle_crestfactor | Signal descriptors | Crest factor of instantaneous turning angle (peak divided by RMS). |
| 87 | inst_angle_formfactor | Signal descriptors | Form factor of instantaneous turning angle (RMS divided by mean absolute value). |
| 88 | inst_angle_pulseindicator | Signal descriptors | Pulse indicator of instantaneous turning angle (peak divided by mean absolute value). |
| 89 | inst_angle_approximate_entropies | Entropy | Approximate entropy of the instantaneous turning-angle series. |
| 90 | inst_angle_attention_entropies | Entropy | Entropy of attention-weight distributions derived from the instantaneous turning-angle series. |
| 91 | inst_angle_permutation_entropies | Entropy | Permutation entropy of the instantaneous turning-angle series. |
| 92 | inst_angle_bubble_entropies | Entropy | Bubble entropy of the instantaneous turning-angle series. |
| 93 | inst_angle_cosine_similarity_entropies | Entropy | Entropy of cosine-similarity distributions between windowed segments of the turning-angle series. |

[**Supplementary video 1.**](https://drive.google.com/file/d/1NsUPxrPGE6-FwGgH5X2cVyH4M9SDjuUo/view?usp=sharing) PCLS collapsing during 37°C incubation

[**Supplementary video 2**](https://drive.google.com/file/d/1llT1z1NPP021h_Fk6j_JAqS9Cwur8gCm/view?usp=sharing)**.** Neutrophil adhesion and migration in PCLS. White arrow pointing to a representative neutrophil probing and migrating in PCLS.
